# Between virus correlations in the outcome of infection across host species: evidence of virus genotype by host species interactions

**DOI:** 10.1101/2021.02.16.431403

**Authors:** Ryan M. Imrie, Katherine E. Roberts, Ben Longdon

## Abstract

Virus host shifts are a major source of outbreaks and emerging infectious diseases, and predicting the outcome of novel host and virus interactions remains a key challenge for virus research. The evolutionary relationships between host species can explain variation in transmission rates, virulence, and virus community composition between hosts, but the potential for different viruses to interact with host species effects has yet to be established. Here, we measure correlations in viral load of four *Cripavirus* isolates across experimental infections of 45 *Drosophilidae* host species. We find positive correlations between every pair of viruses tested, suggesting that broadly susceptible host clades could act as reservoirs and donors for certain types of viruses. Additionally, we find evidence of genotype-by-genotype interactions between viruses and host species, highlighting the importance of both host and virus traits in determining the outcome of virus host shifts. More closely related viruses tended to be more strongly correlated, providing tentative evidence that virus evolutionary relatedness may be a useful proxy for determining the likelihood of novel virus emergence, which warrants further research.

**Impact Summary:** Many new infectious diseases are caused by viruses jumping into novel host species. Estimating the probability that jumps will occur, what the characteristics of new viruses will be, and how they are likely to evolve after jumping to new host species are major challenges. To solve these challenges, we require a detailed understanding of the interactions between different viruses and hosts, or metrics that can capture some of the variation in these interactions. Previous studies have shown that the evolutionary relationships between host species can be used to predict traits of infections in different hosts, including transmission rates and the damage caused by infection. However, the potential for different viruses to influence the patterns of these host species effects has yet to be determined. Here, we use four viruses of insects in experimental infections across 45 different fruit fly host species to begin to answer this question. We find similarities in the patterns of replication and persistence between all four viruses, suggesting susceptible groups of related hosts could act as reservoirs and donors for certain types of virus. However, we also find evidence that different virus genotypes interact in different ways with some host species. Viruses that were more closely related tended to behave in similar ways, and so we suggest that virus evolutionary relatedness may prove to be a useful metric for predicting the traits of novel infections and should be explored further in future studies.

## Introduction

Virus host shifts, where viruses jump to and establish onward transmission in novel host species, are a major source of outbreaks and emerging infectious diseases [1–3]. Many human infections, including HIV, Ebola, and recently SARS-CoV-2, have shifted into humans from other species, and continue to cause significant damage to public health, society, and the global economy [4–7]. Predicting and preventing virus host shifts have consequently become major goals of virus research [8]. Many challenges remain in achieving these goals, including improving our understanding of the host, virus, and ecological factors that influence the outcome of initial cross-species transmission [9,10], and the evolutionary and epidemiological factors that determine which pathogens become established in novel hosts [11].

Several studies have investigated how host evolutionary relatedness can explain variation in the outcome of infection across host species. Greater phylogenetic distance between the natural (donor) and recipient hosts is associated with decreased likelihood of cross-species transmission [12,13] and reduced onward transmission within the novel host species [14]. Phylogenetic distance between hosts also explains variation in virulence after cross-species transmission, which increases when viruses jump between more distantly related hosts [14–16]. Groups of closely related hosts have also been shown to share similar levels of susceptibility to novel viruses, independent of the distance to the natural host [17,18], and harbour similar virus communities [19–21].

Less is known about the interactions that exist between virus genotype and host species in determining the outcome of infection [18]. Within species, genotype-by-genotype interactions between host and virus can be important determinants of the outcome of infection. These interactions alter the rank-order of host susceptibility and so reduce the strength of correlations between viruses [22]. Comparative analyses of fungal pathogens of plants [23] and ectoparasites of mammals [24] have revealed parasite phylogenetic effects, and some evidence exists to suggest similar effects may be found in viruses. Closely related viruses tend to infect the same broad host taxa [25], despite high levels of geographic range overlap between potential hosts [26], suggesting they share similar constraints on their host ranges. Both co-speciation and the preferential host switching of viruses can support this assumption, given that viruses are overwhelmingly likely to encounter other host taxa over the timescales required for speciation. That said, shifts between divergent host species are also common across every virus family [27] and these exceptions include several human zoonoses of major concern [14].

The extent to which closely related viruses share traits has yet to be established. However, inferring the characteristics of viruses from better studied relatives is common and sometimes necessary. This is frequently the case during the early stages of outbreaks, where primary research on new viruses or variants is not available. When SARS-CoV-2 first emerged, its characteristics and epidemiological trajectory were inferred from closely related zoonotic and endemic coronaviruses [28], and from other pandemic respiratory viruses such as influenza A [29]. Comparisons to previous outbreaks were used to parameterise disease models in the 2009 H1N1 pandemic [30,31], the 2014 Ebolavirus outbreak [32], and in forecast models of seasonal influenza [33]. Even for viruses that are not newly emerged, many experimental models of infection rely on surrogates when the virus of interest is unavailable, non-permissive in cell culture or animal models, or requires considerable adaptation to experimental hosts [34,35].

These comparisons assume that the traits of one virus are similar to other, related viruses. However, comparisons between more distantly related viruses, such as bat and canine rabies viruses [44] and diverged lineages of influenza viruses [45–47], found stark differences across larger evolutionary scales. Many examples also exist of small genetic changes having large phenotypic effects in viruses, including single SNP changes altering the host range of canine parvoviruses [36], the vector specificity of Chikungunya virus [37], and the infectivity of naturally occurring Ebolaviruses [38]. Only three amino acid substitutions are required to switch receptor specificity of avian H7N9 influenza from poultry to human cell receptors [39]. Virus evolution is often characterised by high mutation rates and frequent reassortment and recombination [40–42]. This, alongside an incomplete sampling of extant viruses [43], has left many poorly resolved evolutionary relationships between and within existing virus lineages [44]. Given these complications, it remains an open question whether comparisons between related viruses can produce accurate inferences of infection traits.

In this study, we have investigated how patterns of host susceptibility (the ability of a virus to persist and replicate) are correlated between viruses, using experimental infections with four *Cripavirus* isolates (family Dicistroviridae) across a panel of 45 host species of *Drosophilidae*. Three of the viruses are isolates of *Drosophila* C virus (DCV-C, DCV-EB and DCV-M), a well-studied virus isolated from *Drosophila melanogaster* [45]. The fourth virus is the closely related Cricket Paralysis virus (CrPV) which was isolated from Australian field crickets (*Teleogryllus commodus*) and is a widely used model insect pathogen [46–48]. Both DCV and CrPV infect a broad range of insect taxa [49], cause virulent infections in adult flies [17,50], and share similar mechanisms for co-opting the host translation machinery [51]. A major-effect resistance gene called pastrel increases resistance to DCV in *D. melanogaster* [52–54] and has also been shown to provide cross-resistance to CrPV along with another gene, Ubc-E2H [55]. Both DCV and CrPV are targeted by the host antiviral RNAi pathway and each encodes a potent suppressor of antiviral RNAi. However, these suppressors have different functions and target different components of the RNAi pathway [56,57]. DCV and CrPV also differ in their tissue pathology; DCV has been shown to infect gut tissues, causing intestinal obstruction following septic inoculation in *D. melanogaster*, which was not observed in CrPV infection [58]. Although little is known about the differences between DCV isolates, they have been shown to cause similar levels of virulence in *D. melanogaster* and interact differently with the endosymbiont *Wolbachia* [59].

Previous work in this host system has shown that susceptibility to DCV-C varies across host species, with the host phylogeny explaining a large proportion of the variance in both viral load and virulence [17]. The host phylogeny is also an important determinant of the evolution of DCV-C in novel hosts, with evidence that mutations that adapt the virus to one host may also adapt it to closely related host species. This suggests virus genotype could alter the likelihood of host shifts in *Drosophila* [60]. Here, we measure correlations in the ability of four viruses to replicate and persist across host species and provide evidence of both broad similarities in infection outcome and differences consistent with virus by host species interactions.

## Materials & Methods

### Fly Stocks

Flies were taken from laboratory stocks of 45 different species of *Drosophilidae* (S1 Text Table A). Before experiments began all included stocks were confirmed to be negative for infection with DCV and CrPV by quantitative reverse transcription PCR (qRT-PCR, described below). Stocks were maintained in multi-generation *Drosophila* stock bottles (Fisherbrand) at 22°C, in a 12-hour light-dark cycle. Each bottle contained 50ml of one of four varieties of food media (S1 Text Supplementary Methods).

### Host Phylogeny

The method used to infer the host phylogeny has been described in detail elsewhere [17]. Briefly, publicly available sequences of the 28S, *Adh, Amyrel, COI, COII, RpL32, and SOD* genes were collected from Genbank (see https://doi.org/10.6084/m9.figshare.13079366.v1 for a full breakdown of genes and accessions by species). Gene sequences were aligned in Geneious v9.1.8 (https://www.geneious.com) using a progressive pairwise global alignment algorithm with free end gaps and a 70% similarity IUB cost matrix. Gap open penalties, gap extension penalties, and refinement iterations were kept as default.

Phylogenetic reconstruction was performed using BEAST v1.10.4 [61] as the subsequent phylogenetic mixed model (see below) requires a tree with the same root-tip distances for all taxa. Genes were partitioned into separate ribosomal (28S), mitochondrial (*COI, COII*), and nuclear (*Adh, Amyrel, RpL32, SOD*) groups. The mitochondrial and nuclear groups were further partitioned into groups for codon position 1+2 and codon position 3, with unlinked substitution rates and base frequencies across codon positions. Each group was fitted to separate relaxed uncorrelated lognormal molecular clock models using random starting trees and 4-category gamma-distributed HKY substitution models. The BEAST analysis was run twice, with 1 billion MCMC generations sampled every 100,000 iterations, using a birth-death process tree-shape prior. Model trace files were evaluated for chain convergence, sampling, and autocorrelation using Tracer v1.7.1 [62]. A maximum clade credibility tree was inferred from the posterior sample with a 10% burn-in. The reconstructed tree was visualised using ggtree v2.0.4 [63].

### Virus Isolates

Virus stocks were kindly provided by Julien Martinez (DCV isolates) [59], and Valérie Dorey and Maria Carla Saleh (CrPV) [57]. DCV-C, DCV-EB and DCV-M were originally isolated from fly stocks with origins in three separate continents; DCV-C and DCV-EB were isolated from lab stocks established by wild capture in Charolles, France and Ellis Beach, Australia respectively, while DCV-M was isolated directly from wild flies in Marrakesh, Morocco [45]. The CrPV isolate was collected from *Teleogryllus commodus* in Victoria, Australia [64]. Virus stocks were diluted in Ringers solution [65] to equalise the relative amounts of viral RNA and checked for contamination with CrPV (DCV isolates) and DCV (CrPV isolate) by qRT-PCR as described below.

### Virus phylogeny

Full genome sequences for DCV-C (*MK645242*), DCV-EB (*MK645239*), DCV-M (*MK645243*), and CrPV (*NC_003924*) were retrieved from the NCBI Nucleotide database. Annotations of ORFs for the replicase polyprotein (CrPV: *Q9IJX4*, DCV: *O36966*) and structural polyprotein (CrPV: *P13418*, DCV: *O36967*) were collected from the UniProtKB database and used to separate the coding and non-coding regions of each virus. ORF sequences were concatenated and aligned using the Geneious progressive pairwise translation alignment algorithm with a Blosum50 cost matrix and default parameters. Alignments were manually checked for quality and sequences aligning to CrPV ORF1 nucleotides 1-387 and 2704-2728 were removed due to the presence of large indels.

Phylogenetic reconstruction was performed using BEAST v1.10.4 with translated ORF sequences fitted to an uncorrelated relaxed lognormal molecular clock model using a speciation birth-death process tree-shape prior. A Blosum62 substitution model [66] with a gamma distribution of rate variation with four categories and a proportion of invariable sites was used. The model was run for 10 million MCMC generations sampled every 1,000 iterations and evaluated in Tracer v1.7.1 as above, and a maximum clade credibility tree inferred with a 10% burn-in.

### Inoculation

Before inoculation, 0-1 day old male flies were kept in vials containing cornmeal media (S1 Text Supplementary Methods), and were transferred to fresh media every 2 days for one week. Vials contained between 5 and 20 flies (mean = 14.5) and were kept at 22°C at 70% relative humidity in a 12-hour light-dark cycle. Flies were inoculated at 7-8 days old under CO_2_ anaesthesia via septic pin prick with 12.5μm diameter stainless steel needles (Fine Science Tools, CA, USA). These needles were bent approximately 250μm from the end to provide a depth stop and dipped in virus solution before being pricked into the pleural suture of each fly. Inoculation by this method has been shown to follow the same course as oral infection but is less stochastic [67]. Inoculated flies were then snap frozen immediately in liquid nitrogen, providing a 0 days-post-infection (dpi) timepoint, or maintained in cornmeal vials for a further 2 days ± 3 hours before freezing, providing a 2 dpi time point. Within replicate blocks the 0 and 2 dpi vials for each virus were inoculated on the same day, and together constituted one biological replicate. We aimed to collect three biological replicates for each species and virus combination, with the order of species, vial (0 or 2 dpi), and virus randomised for each replicate block.

### Measuring Change in Viral Load

To measure the change in viral load between 0 and 2 dpi, total RNA was extracted from flies homogenised in Trizol (Invitrogen, supplied by ThermoFisher) using chloroform-isoproponyl extraction, and reverse transcribed using Promega GoScript reverse transcriptase (Sigma) with random hexamer primers. qRT-PCR was carried out on 1:10 diluted cDNA on an Applied Biosystems StepOnePlus system using Sensifast Hi-Rox Sybr kit (Bioline). Cycle conditions were as follows: initial denaturation at 95°C for 120 seconds, then 40 cycles of 95°C for 5 seconds, and 60°C for 30 seconds.

DCV isolates were measured using the same primer pair (Forward: 5’-GACACTGCCTTTGATTAG-3’, Reverse: 5’-CCCTCTGGGAACTAAATG-3’) which targeted a conserved location and had similarly high efficiencies across all isolates. For CrPV the following primers were used: Forward: 5’-TTGGCGTGGTAGTATGCGTAT-3’, Reverse: 5’-TGTTCCGTCCTGCGTCTC-3’. *RpL32* housekeeping gene primers varied by species (S1 Text Tables B & C). For each sample, two technical replicates were performed for each amplicon (viral and *RpL32*).

Between-plate variation in C_T_ values was estimated and corrected for using a linear model with plate ID and biological replicate ID as parameters, as described elsewhere [68,69]. Mean viral C_T_ values from technical replicate pairs were normalised to *RpL32* and converted to fold-change in viral load using the 2^-ΔΔCT^ method, where ΔC_T_ = C_T:Virus_ – C_T:Rpl32_, and ΔΔC_T_ = ΔC_T:day0_ - ΔC_T:day2_.

Amplification of the correct products was verified by melt curve analysis. Repeated failure to amplify product, the presence of melt curve contaminants, or departures from the melt curve peaks of positive samples (± 1.5°C for viral amplicons, ± 3°C for Rpl32) in either the 0 or 2 dpi samples were used as exclusion criteria for biological replicates. In total, of the 180 unique combinations of host species and virus measured, 3 biological replicates were obtained for 161 combinations, 2 replicates for 18 combinations, and 1 replicate for 1 combination (*Drosophila* virilis, CrPV).

### Statistical Analysis

Phylogenetic generalised linear mixed models were used to investigate the effects of host relatedness on viral load, and to examine correlations between the different virus isolates. Multivariate models were fitted using the R package MCMCglmm [70] with the viral load of each virus isolate as the response variable. The structures of the models were as follows:

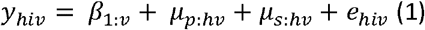

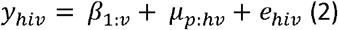

In these models, *y*_*hiv*_ is the change in viral load for virus *v* in the *i*^*th*^ biological replicate of host species *h*. The fixed effect *β*_*1*_ represents the intercepts for each virus isolate, the random effect *µ*_*p*_ represents the effects of the host phylogeny assuming a Brownian motion model of evolution, and *e* represents the model residuals. Model (1) also includes a species-specific random effect that is independent of the host phylogeny (*µ*_*s:hv*_). This explicitly estimates the non-phylogenetic component of between-species variance and allows the proportion of variance explained by the host phylogeny to be calculated. *µ*_*s:hv*_ was removed from model (2) as model (1) struggled to separate the phylogenetic and species-specific traits. Wing size, measured as the length of the IV longitudinal vein from the tip of the proximal segment to the join of the distal segment with vein V [71], provided a proxy for body size [72] and was included in a further model as a fixed effect (*wingsize*_*h*_.*β*_*2*_).

Within each of these models the random effects and residuals were assumed to follow a multivariate normal distribution with a centred mean of zero and a covariance structure of V_p_⊗ A for the phylogenetic effects, V_s_⊗ I for species-specific effects, and V_e_⊗ I residuals, where ⊗ represents the Kronecker product. A represents the host phylogenetic relatedness matrix, I an identity matrix and V represents 4×4 covariance matrices describing the between-species variances and covariances of changes in viral load for the different viruses. Specifically, the matrices *V*_*p*_ and *V*_*s*_ describe the phylogenetic and non-phylogenetic between-species variances in viral load for each virus and the covariances between them, while the residual covariance matrix *V*_*e*_ describes within-species variance that includes both true within-species effects and measurement errors. Since each biological replicate was tested with a single virus isolate, the covariances of *V*_*e*_ cannot be estimated and were set to zero.

Models were run for 13 million MCMC generations, sampled every 5,000 iterations with a burn-in of 3 million generations. Parameter expanded priors were placed on the covariance matrices, resulting in multivariate F distributions with marginal variance distributions scaled by 1,000. Inverse-gamma priors were placed on the residual variances, with a shape and scale equal to 0.002. To ensure the model outputs were robust to changes in prior distribution, models were also fitted with flat and inverse-Wishart priors, which gave qualitatively similar results.

The proportion of the between species variance that can be explained by the phylogeny was calculated from model (1) using the equation *v*_*p*_ */(v*_*p*_ *+ v*_*s*_), where *v*_*p*_ and *v*_*s*_ represent the phylogenetic and species-specific components of between-species variance [73], and is equivalent to phylogenetic heritability or Pagel’s lambda [74,75]. The repeatability of viral load measurements was calculated from model (2) as *v*_*p*_*/(v*_*p*_ *+ v*_*e*_), where *V*_*e*_ is the residual variance of the model [76]. Inter-specific correlations in viral load were calculated from model (2) V_p_ matrix as *cov*_*x,y*_*/v(var*_*x*_ *+ var*_*y*_). If correlations between viruses are close to one (with no change in the variance whilst the means

remain constant) it would suggest there are no host species-by-virus interactions [22]. Parameter estimates reported are means of the posterior density, and 95% credible intervals (95% CI) were taken to be the 95% highest posterior density intervals.

The data files and R scripts used in this study are available in an online repository: https://doi.org/10.6084/m9.figshare.13750711.v1.

## Results

### Change in viral load is a repeatable trait among host species

To investigate similarities between related viruses in the outcome of infection across host species, as well as the potential for virus genotype to interact with host species effects, we experimentally infected 45 species of *Drosophilidae* with four virus isolates: DCV-C, DCV-EB, DCV-M, and CrPV. The DCV isolates formed a distinct clade (>93% genome and ORF amino acid identity, with 265-556 SNPs between isolates), with the closest relationship between DCV-C and DCV-EB. CrPV formed an outgroup to the DCV isolates (57-59% identity, with over 4000 SNPs between CrPV and each DCV isolate, Fig. 1, Table 1). In total, 15,657 flies were inoculated, and the change in viral load after two days of infection was determined by qRT-PCR (Fig. 2). The mean viral load within host species ranged from an approximately 2.7-billion-fold increase in *Drosophila* persimilis infected with DCV-M to a 2.5-fold decrease in *Zaprionus tuberculatus* infected with DCV-C. Viral loads across host species tended to be higher for the DCV isolates, with a mean fold-increase of roughly 11,000 - 19,000, and lower for CrPV, with a mean fold-increase of roughly 1,600.

**Figure 1.**
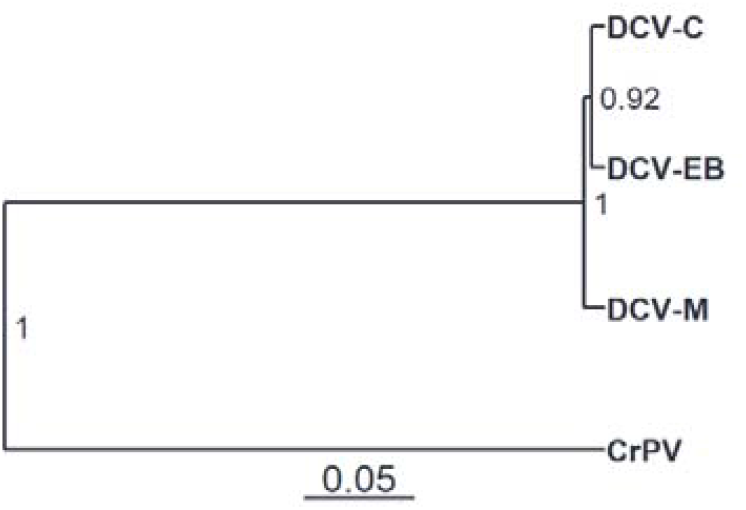
Phylogeny of virus isolates. Evolutionary relationships estimated from open reading frame (ORF) amino acid sequences presented in a midpoint-rooted tree. Node labels represent the posterior probabilities of each clade, and the scale bar represents amino acid substitutions per site.

**Table 1:** Virus isolate sequence similarity. Percentage sequence identity was calculated from multiple-alignment of whole genome nucleotides (white) or concatenated amino acid sequences of ORFs 1 & 2 (grey). Approximately 92 SNPs and 28 amino acid substitutions exist for every 1% of sequence divergence.

|  | DCV-C | DCV-EB | DCV-M | CrPV |
| --- | --- | --- | --- | --- |
| DCV-C |  | 97.10% | 94.00% | 57.40% |
| DCV-EB | 98.81% |  | 93.90% | 57.30% |
| DCV-M | 98.30% | 98.20% |  | 57.40% |
| CrPV | 59.00% | 58.70% | 58.70% |  |

**Figure 2.**
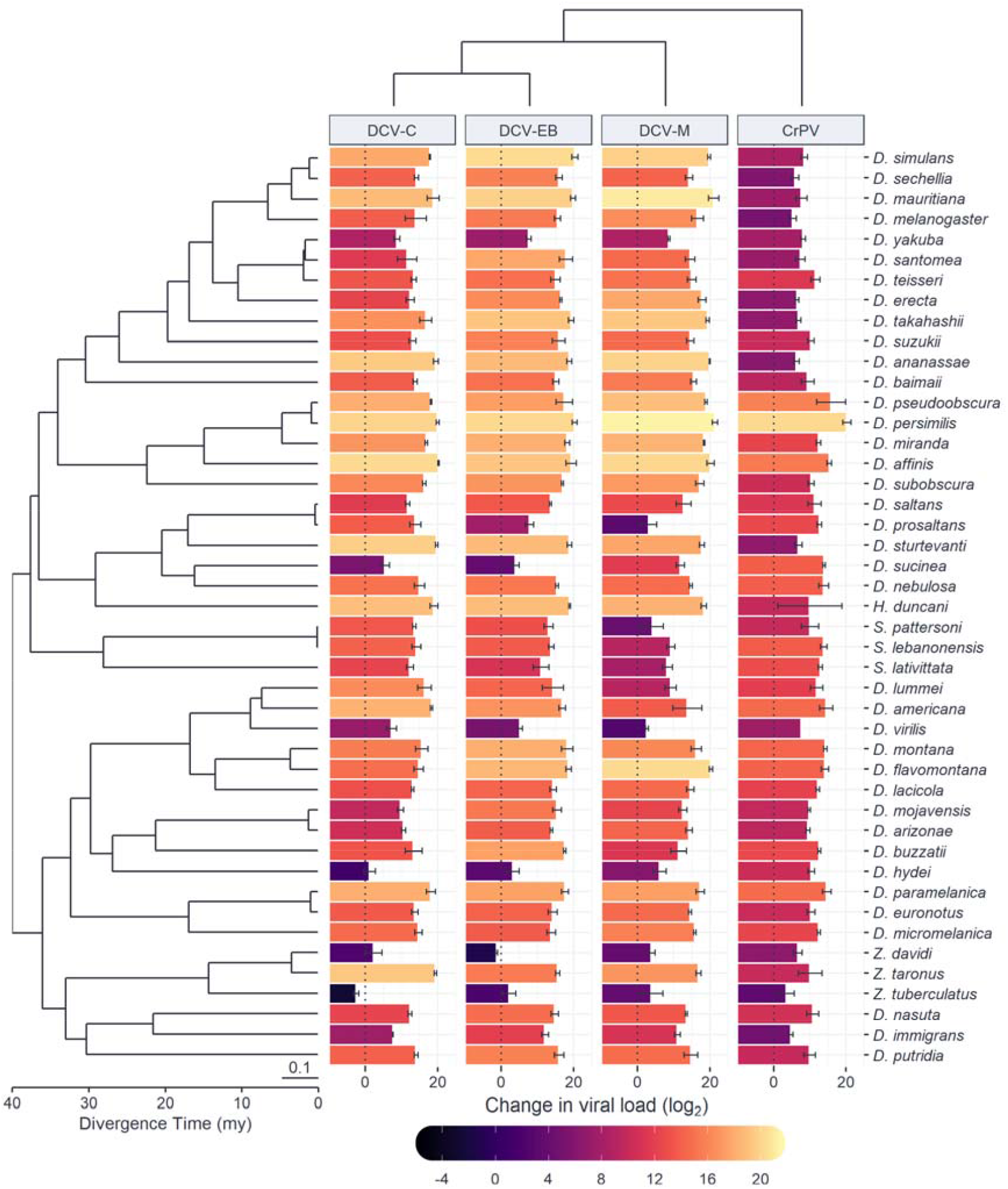
Change in viral load across a diverse panel of *Drosophilidae* host species for different virus isolates. Bar height and colour show the mean change in viral load at 2 dpi on a log_2_ scale, with error bars representing the standard error of the mean. The phylogeny of *Drosophilidae* hosts is presented on the left, with the scale bar representing the number of nucleotide substitutions per site and scale axis representing the approximate age since divergence in millions of years (my) based on estimates from [77,78]. The virus cladogram, presented at the top, is based on the evolutionary relationships shown in Fig. 1.

Phylogenetic generalised linear mixed models were fitted to the data to determine the proportion of variation in viral load explained by the host phylogeny (Table 2). The phylogeny explained 79% of the variation in viral load for CrPV but only 9-21% of the variation for the DCV isolates, with wide credible intervals on all the DCV estimates. This was due to the model struggling to separate phylogenetic and species-specific effects for these viruses. The repeatability of viral load across host species was high for both CrPV (0.66) and the DCV isolates (0.92-0.96), with the between-species phylogenetic component (*v*_*p*_) explaining a high proportion of the variation in viral load with little within-species variation or measurement error (*v*_*p*_). We found no significant effect of wing length (a proxy for host body size) on viral load for any of the included viruses, with all estimates having credible intervals overlapping zero.

**Table 2:** Estimates of mean change in viral load, repeatability, and the proportion of variation explained by the host phylogeny. Estimates of the mean change in viral load and repeatability are taken from model (2), while estimates of the variation explained by the host phylogeny are taken from model (1).

| Virus | Mean change in viral load ($\log_2$) | Repeatability | Variance explained by phylogeny |
| --- | --- | --- | --- |
| DCV-C | 13.50 (95% CI: 11.17, 15.89) | 0.96 (95% CI: 0.93, 0.98) | 0.11 (95% CI: 0, 0.35) |
| DCV-EB | 14.22 (95% CI: 11.42, 16.76) | 0.96 (95% CI: 0.93, 0.98) | 0.09 (95% CI: 0, 0.32) |
| DCV-M | 13.63 (95% CI: 10.52, 16.59) | 0.92 (95% CI: 0.87, 0.96) | 0.23 (95% CI: 0, 0.51) |
| CrPV | 10.66 (95% CI: 8.59, 12.66) | 0.66 (95% CI: 0.46, 0.83) | 0.79 (95% CI: 0.50, 1.00) |

### Correlations between viruses are consistent with virus by host species interactions

Inter-specific correlations in viral load between viruses were then estimated from the variance-covariance matrices of model (2) (Fig. 3A). We found strong positive correlations between the DCV isolates (r > 0.93), with the strongest correlation between DCV-C and DCV-EB (r = 0.97). Correlations between DCV isolates and the more distantly related CrPV were positive (r = 0.52-0.59) but weaker than the correlations between the DCV isolates. The fact the DCV:CrPV correlations (and their 95% CI’s) are not close to one is consistent with virus genotype by host species interactions on viral load [22]. This is further demonstrated by the notable differences in the rank order of host species susceptibility for each virus (Fig. 3B), equivalent to a crossing over of reaction norms for the susceptibility of host species between virus genotypes [79].

**Figure 3.**
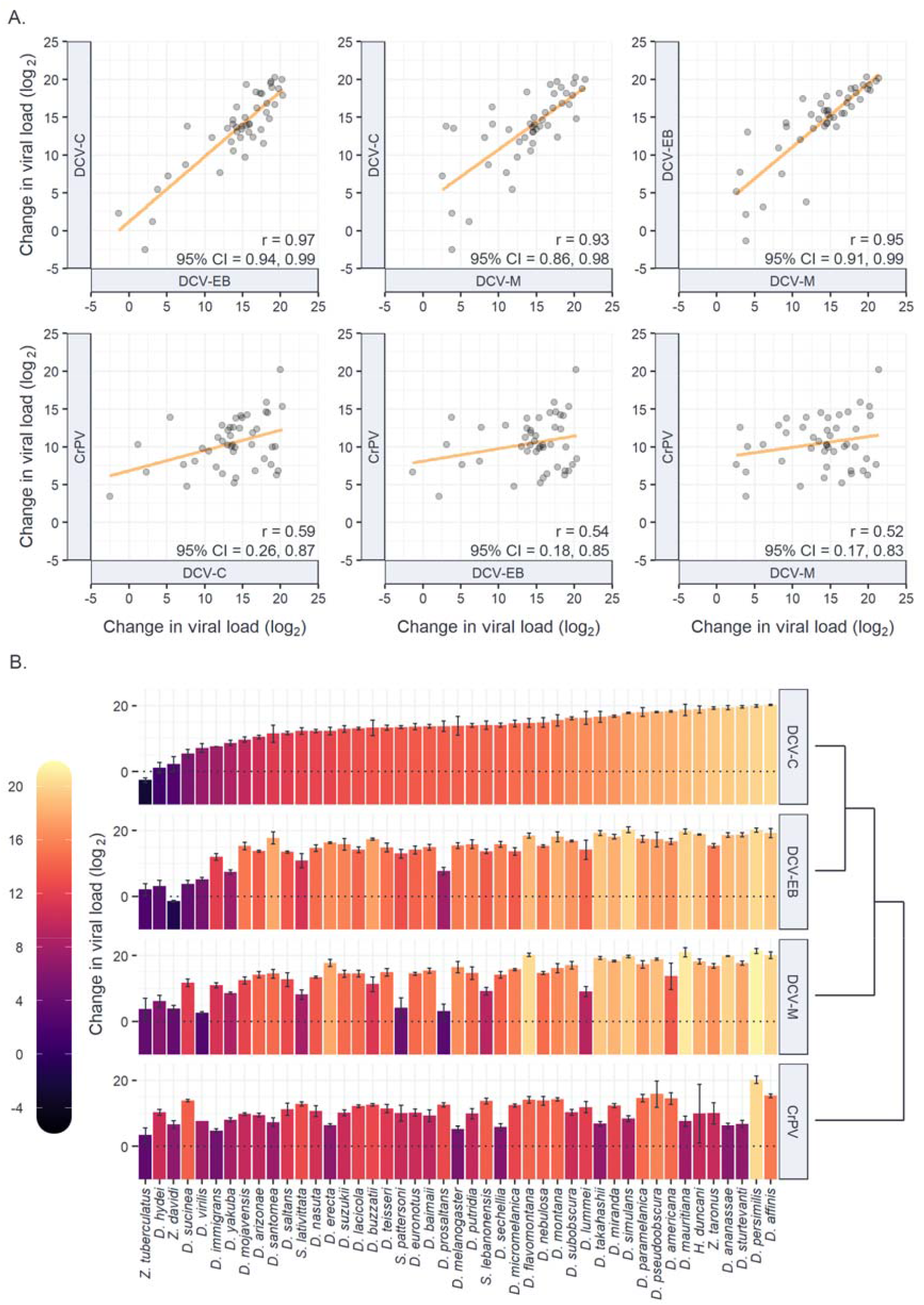
Similarities in infection outcome across host species and interactions between virus genotype and host species. A) Correlations in viral load between virus isolates. Individual points represent the mean change in viral load after 2 dpi for each host species on a log_2_ scale, and trendlines have been added from a univariate least-squares linear model for illustrative purposes. Correlations (r) are the total inter-specific correlations and 95% CIs from the output of model (2). B) Differences in the rank-order of host species susceptibility between virus isolates. Bar height and colour show the mean change in viral load at 2 dpi on a log_2_ scale, with error bars representing the standard error of the mean. The order of species along the x-axis has been sorted in ascending order of viral load during infection with DCV-C. Deviations from this rank-order of host species susceptibility for other viruses is indicative of crossing reaction norms and interactions between virus genotype and host species. The virus cladogram is based on the evolutionary relationships shown in Fig. 1.

DCV-C was more strongly correlated to DCV-EB than to DCV-M (Δr = 0.04, 95% CI = 0.00, 0.09), and more strongly correlated to DCV-M than to CrPV (Δr = 0.40, 95% CI = 0.18, 0.82), consistent with an increase in the strength of correlation between viruses with closer evolutionary relatedness. Point estimates imply a similar pattern for DCV-EB, but the evidence for a stronger correlation with DCV-C than DCV-M was not well supported (S1 Text Table D).

## Discussion

Closely related host species present similar environments to novel viruses [73,80], and so tend to share similar levels of susceptibility to a given virus [12–18]. Likewise, closely related viruses are often assumed to share characteristics that make their host interactions, transmission, and evolutionary trajectories comparable [28–35]. Here, we measured the strength of correlations in viral load between four *Cripavirus* isolates across 45 host species of *Drosophilidae*, to look for similarities between related viruses as well as evidence of virus genotype by host species interactions on the outcome of infection. We found positive correlations between every pair of viruses tested, indicating broad similarities in the outcome of infection across host species, but also evidence for interactions between virus genotype and host species with changes in the rank order of host species susceptibility between the different viruses (Fig. 3). This highlights the importance of considering both host and virus traits in understanding the outcomes of virus host shifts.

The strong positive correlations between DCV isolates are likely due to relatively high levels of sequence conservation resulting in only small differences in their ability to infect different host species. However, in other viruses a small number of mutations have been shown to allow successful infections in novel hosts [36,39]. We find a few instances of such effects here. For example, in *Zaprionus davidi*, DCV-EB shows a decline in viral load, suggesting it is failing to replicate and persist in this host species, whereas the other isolates show an increase in viral load in the same host. Similarly, *Scaptodrosophila pattersoni* is amongst the least susceptible to DCV-M but has relatively high viral loads for the other virus isolates. A greater number of these effects can be seen when comparing hosts infected with DCV isolates to those infected with CrPV, where multiple species have markedly different susceptibilities depending on the virus infecting them. For example, both *Drosophila* ananassae and *Drosophila* strutevanti are within the five most susceptible species to DCV-C, but also the eight least susceptible to CrPV. The weaker correlations that exist between DCV and CrPV may be due to interactions with different host traits that vary in their patterns across the host phylogeny. CrPV and DCV are known to have distinct methods of suppression of the host antiviral RNAi pathway [56,57] and cause pathology in different tissues [62]. Additionally, their relatively high levels of sequence divergence (57-59% identity) may have resulted in changes in the ability of each virus to bind to host cell receptors, utilise host replication machinery, or avoid host immune defences [81].

The existence of correlations between viruses suggests that host susceptibility is not specific to individual viruses and that certain host clades may be broadly susceptible to infection. Host species that are permissive to multiple virus genotypes may allow for the persistence of increased genetic diversity in the virus population, allowing viruses to generate and maintain mutations that make them more likely to emerge in novel host species [82,83]. They also have the potential to act as “mixing vessels”, providing increased opportunities for virus reassortment and recombination [84], which has been proposed as a possible route for several viruses to acquire pandemic potential [85,86]. Broadly susceptible host clades may therefore act as common reservoirs and donors of emerging infectious diseases and identifying them in relevant systems could inform control and prevention strategies [87].

The differences in correlation strength between pairs of viruses tended to follow differences in their evolutionary divergence, such that more closely related pairs of viruses were more strongly correlated in the outcome of infection across host species. This provides some tentative evidence that the ability of a virus to infect a novel host may be inferred based on its evolutionary relatedness to other viruses, although we lack the sample size or diversity of viruses needed to test this conclusively. Numerous examples exist where a small number of genetic changes in viruses cause large phenotypic differences [36–39], which would be exceptions to any link between correlation strength and evolutionary relatedness [87]. Nevertheless, virus phylogenetic effects may still prove to be a useful proxy for determining the likelihood of novel virus emergence. Further work is now needed to expand the findings of this study to broader groups of viruses, and to test the importance of the virus phylogeny in determining the potential outcomes of virus host shifts.

## Acknowledgements

We would like to thank Julien Martinez, Valérie Dorey, and Maria Carla Saleh for kindly providing us with DCV and CrPV virus isolates, Jarrod Hadfield for his advice on MCMCglmm model fitting, and Joanne Lello for useful discussions. Ryan M. Imrie is supported by a studentship funded by the Natural Environment Research Council (NERC) GW4+ Doctoral Training Partnership and the College of Life and Environmental Sciences, University of Exeter. Ben Longdon and Katherine E. Roberts are supported by a Sir Henry Dale Fellowship jointly funded by the Wellcome Trust and the Royal Society (Grant Number 109356/Z/15/Z). For the purpose of Open Access, the author has applied a CC BY public copyright licence to any Author Accepted Manuscript version arising from this submission.

