## Supplementary Text 1 for "Between virus correlations in the outcome of infection across host species: evidence of virus genotype by host species interactions": S1_Text.html


### S1 Text

#### **S1 Text Supplementary Methods.**

##### **Drosophila Media Recipes.**

To prepare media, ingredients (excluding nipagin and proprionic acid) are mixed together and brought to boil, then allowed to cool to 80°C before adding preservatives and pouring into bottles. Media is allowed to solidify overnight at room temperature before use.

**Banana:**

| Ingredient | Quantity |
| --- | --- |
| Water | 1,000ml |
| Agar | 10g |
| Yeast | 30g |
| Bananas^\*^ | 150g |
| Malt Extract | 30g |
| Molasses | 50g |
| Nipagin | 25ml |
|  |  |
| --- | --- |
| ^*\**^  Blended in water before boiling. | |

  
 **Cornmeal:**

| Ingredient | Quantity |
| --- | --- |
| Water | 1,000ml |
| Agar | 11g |
| Yeast | 19g |
| Cornmeal | 88g |
| Dextrose | 88g |
| Nipagin | 29ml |

**Malt:**

| Ingredient | Quantity |
| --- | --- |
| Water | 1,000ml |
| Agar | 10g |
| Yeast | 20g |
| Semolina | 60g |
| Malt Extract | 80g |
| Nipagin | 14ml |
| Propionic Acid | 5ml |

**Propionic:**

| Ingredient | Quantity |
| --- | --- |
| Water | 1,000ml |
| Agar | 10g |
| Yeast | 20g |
| Cornmeal | 70g |
| Soya Flour | 10g |
| Malt Extract | 80g |
| Molasses | 22g |
| Nipagin | 14ml |
| Propionic Acid | 6.2ml |

#### **S1 Text Supplementary Tables**

##### **S1 Text Table A.** Full list of Drosophila host species used in this study, including wingspan data used as a proxy for body size and the media type used to maintain each stock population. Recipes for each media type can be found below.

| Species | Genus | Wingsize (mm) | Media |
| --- | --- | --- | --- |
| D. affinis | Drosophila | 1.803 | Malt |
| D. americana | Drosophila | 2.045 | Malt |
| D. ananassae | Drosophila | 1.493 | Cornmeal |
| D. arizonae | Drosophila | 1.548 | Banana |
| D. baimaii | Drosophila | 1.561 | Cornmeal |
| D. buzzati | Drosophila | 1.902 | Malt |
| D. erecta | Drosophila | 1.581 | Malt + yeast |
| D. euronotus | Drosophila | 2.222 | Cornmeal |
| D. flavomontana | Drosophila | 2.192 | Malt + yeast |
| D. hydei | Drosophila | 2.182 | Cornmeal |
| D. immigrans | Drosophila | 2.153 | Malt + yeast |
| D. lacicola | Drosophila | 2.268 | Malt |
| D. lummei | Drosophila | 2.558 | Malt + yeast |
| D. mauritiana | Drosophila | 1.507 | Proprionic |
| D. melanogaster | Drosophila | 1.716 | Cornmeal |
| D. micromelanica | Drosophila | 1.895 | Cornmeal |
| D. miranda | Drosophila | 2.395 | Cornmeal |
| D. mojavensis | Drosophila | 1.650 | Banana |
| D. montana | Drosophila | 2.706 | Malt + yeast |
| D. nasuta | Drosophila | 1.917 | Cornmeal |
| D. nebulosa | Drosophila | 1.826 | Cornmeal |
| D. paramelanica | Drosophila | 1.946 | Cornmeal |
| D. persimilis | Drosophila | 2.013 | Malt |
| D. prosaltans | Drosophila | 1.699 | Proprionic |
| D. pseudoobscura | Drosophila | 1.863 | Malt |
| D. putridia | Drosophila | 1.639 | Proprionic |
| D. saltans | Drosophila | 1.600 | Proprionic |
| D. santomea | Drosophila | 1.489 | Cornmeal |
| D. sechellia | Drosophila | 1.424 | Proprionic |
| D. simulans | Drosophila | 1.484 | Cornmeal |
| D. sturtevanti | Drosophila | 1.779 | Cornmeal |
| D. subobscura | Drosophila | 2.056 | Cornmeal |
| D. sucinea | Drosophila | 1.932 | Cornmeal |
| D. suzukii | Drosophila | 2.100 | Cornmeal |
| D. takahashii | Drosophila | 1.559 | Cornmeal |
| D. teissieri | Drosophila | 1.463 | Cornmeal |
| D. virilis | Drosophila | 2.253 | Proprionic |
| D. yakuba | Drosophila | 1.307 | Cornmeal |
| H. duncani | Hirtodrosophila | 1.969 | Proprionic |
| S. lativittata | Scaptodrosophila | 1.851 | Banana |
| S. lebanonensis | Scaptodrosophila | 2.053 | Proprionic |
| S. pattersoni | Scaptodrosophila | 2.023 | Banana |
| Z. davidi | Zaprionous | 1.911 | Banana |
| Z. taronus | Zaprionous | 2.131 | Banana |
| Z. tuberculatus | Zaprionous | 1.914 | Banana |
|  |  |  |  |
| --- | --- | --- | --- |
| Cornmeal and proprionic media are dusted with yeast before use. Malt and banana media are left bare unless otherwise stated. | | | |

##### **S1 Text Table B.** *RPL32* primer sequences used to amplify the host housekeeping gene. Primers were designed to amplify across an intron boundary to ensure only cDNA sequences were amplified. Different primer combinations were used for different host species (S1 Text Table C) to account for SNPs in the primer binding sites.

|  | Name | Sequence |
| --- | --- | --- |
| Forward | RpL32\_qPCR\_F-a | TGCCAAGTTGTCGCACAAATGG |
|  | RpL32\_qPCR\_F-b | TGCTAAGTTGTCGCACAAATGG |
|  | RpL32\_qPCR\_F-c | TGCCAAGCTGTCGCACAAATGG |
|  | RpL32\_qPCR\_F-d | TGCTAAGCTGTCGCACAAATGG |
|  | RpL32\_qPCR\_F-e | TGCGAAGTTGTCGCACAAATGG |
|  | RpL32\_qPCR\_F-f | TGCGAAGCTGTCGCACAAATGG |
| Reverse | RpL32\_qPCR\_R-a | TGCGCTTGTTGGAACCGTAAC |
|  | RpL32\_qPCR\_R-b | TGCGCTTGTTGGATCCGTAAC |
|  | RpL32\_qPCR\_R-c | TGCGCTTGTTGGAACCATAAC |
|  | RpL32\_qPCR\_R-d | TGCGCTTGTTGGAGCCGTAAC |
|  | RpL32\_qPCR\_R-e | TGCGCTTGTTAGAACCGTAAC |
|  | RpL32\_qPCR\_R-f | TACGCTTGTTGGAACCGTAAC |
|  | RpL32\_qPCR\_R-g | TGCGCTTGTTGGAACCGTAGC |
|  | RpL32\_qPCR\_R-h | TGCGCTTGTTCGATCCGTAAC |
|  | RpL32\_qPCR\_R-i | TGCGCTTGTTGGAGCCATAAC |
|  | RpL32\_qPCR\_R-j | TGCGCTTGTTTGATCCGTAAC |
|  | RpL32\_qPCR\_R-k | TGCGCTTGTTTGAACCATAAC |
|  | RpL32\_qPCR\_R-l | TACGCTTGTTGGAACCATAAC |
|  | RpL32\_qPCR\_R-m | TACGCTTGTTGGAGCCGTAAC |
|  | RpL32\_qPCR\_R-n | TGCGCTGGTTGGAACCATAAC |
|  | RpL32\_qPCR\_R-o | TGAGCTTGTTCGATCCGTAAC |
|  | RpL32\_qPCR\_R-p | TACGCTTGTTGGAGCCATAAC |
|  | RpL32\_qPCR\_R-q | TGAGCTTGTTTGATCCGTAAC |
|  | RpL32\_qPCR\_R-r | TAAGCTTGTTGGATCCGTAGC |
|  | RpL32\_qPCR\_R-s | TCAGCTTGTTGGATCCATAGC |

##### **S1 Text Table C.** *RPL32* primer combinations used for each host species.

| Species | Forward | Reverse |
| --- | --- | --- |
| D. affinis | F-a | R-i |
| D. americana | F-c | R-a |
| D. ananassae | F-f | R-a |
| D. arizonae | F-a | R-a |
| D. baimaii | F-a | R-r |
| D. buzzati | F-a | R-e |
| D. erecta | F-d | R-h |
| D. euronotus | F-a | R-g |
| D. flavomontana | F-c | R-a |
| D. hydei | F-a | R-a |
| D. immigrans | F-b | R-p |
| D. lacicola | F-c | R-a |
| D. lummei | F-c | R-a |
| D. mauritiana | F-d | R-h |
| D. melanogaster | F-d | R-h |
| D. micromelanica | F-a | R-g |
| D. miranda | F-a | R-d |
| D. mojavensis | F-a | R-a |
| D. montana | F-c | R-a |
| D. nasuta | F-b | R-f |
| D. nebulosa | F-b | R-c |
| D. paramelanica | F-a | R-g |
| D. persimilis | F-a | R-b |
| D. prosaltans | F-a | R-n |
| D. pseudoobscura | F-a | R-m |
| D. putridia | F-d | R-q |
| D. saltans | F-a | R-n |
| D. santomea | F-a | R-n |
| D. sechellia | F-d | R-h |
| D. simulans | F-d | R-h |
| D. sturtevanti | F-a | R-l |
| D. subobscura | F-a | R-i |
| D. sucinea | F-b | R-k |
| D. suzukii | F-d | R-o |
| D. takahashii | F-d | R-o |
| D. teissieri | F-d | R-h |
| D. virilis | F-c | R-a |
| D. yakuba | F-d | R-h |
| H. duncani | F-f | R-c |
| S. lativittata | F-a | R-m |
| S. lebanonensis | F-d | R-h |
| S. pattersoni | F-a | R-m |
| Z. davidi | F-a | R-c |
| Z. taronus | F-a | R-c |
| Z. tuberculatus | F-a | R-c |

##### **S1 Text Table D.** Estimates of the differences between correlations in viral load (Δr). The differences presented have all been calculated as column minus row. Evidence for significant differences between correlations are shown as the mean and 95% CIs for the posterior distribution of the differences in correlations (white) and as P_MCMC_ (grey). P_MCMC_ corresponds to 2 \* P_min_, where P_min_ is either the probability of iterations in the posterior density being positive or negative, whichever is smaller. Significant differences (95% CIs not crossing zero, P_MCMC_ < 0.05) are highlighted in bold.

|  | DCV-C : DCV-EB | DCV-C : DCV-M | DCV-EB : DCV-M | CrPV : DCV-C | CrPV : DCV-EB | CrPV : DCV-M |
| --- | --- | --- | --- | --- | --- | --- |
| DCV-C : DCV-EB |  | -0.04 (95% CI = -0.09, 0.00) | -0.02 (95% CI = -0.07, 0.02) | -0.37 (95% CI = -0.70, -0.13) | -0.43 (95% CI = -0.78, -0.16) | -0.45 (95% CI = -0.82, -0.18) |
| DCV-C : DCV-M | 0.04 (p = 0.03) |  | 0.03 (95% CI = -0.01, 0.08) | -0.33 (95% CI = -0.68, -0.10) | -0.38 (95% CI = -0.77, -0.13) | -0.40 (95% CI = -0.77, -0.16) |
| DCV-EB : DCV-M | 0.02 (p = 0.47) | -0.03 (p = 0.17) |  | -0.36 (95% CI = -0.69, -0.11) | -0.41 (95% CI = -0.73, -0.09) | -0.43 (95% CI = -0.78, -0.16) |
| CrPV : DCV-C | 0.37 (p < 0.001) | 0.33 (p < 0.001) | 0.36 (p < 0.001) |  | -0.05 (95% CI = -0.14, 0.03) | -0.07 (95% CI = -0.23, 0.06) |
| CrPV : DCV-EB | 0.43 (p < 0.001) | 0.38 (p < 0.001) | 0.41 (p < 0.001) | 0.05 (p = 0.24) |  | -0.02 (95% CI = -0.17, 0.11) |
| CrPV : DCV-M | 0.45 (p < 0.001) | 0.40 (p < 0.001) | 0.43 (p < 0.001) | 0.07 (p = 0.32) | 0.02 (p = 0.76) |  |
